# Hnf4a mediates microbial control of intestinal gene expression and inflammation

**DOI:** 10.1101/097055

**Authors:** James M. Davison, Colin R. Lickwar, Lingyun Song, Ghislain Breton, Gregory E. Crawford, John F. Rawls

**Affiliations:** Department of Molecular Genetics and Microbiology, Center for the Genomics of Microbial Systems, Duke University, Durham, North Carolina 27710, USA; Department of Cell Biology and Physiology, University of North Carolina, Chapel Hill, North Carolina 27599, USA; Department of Pediatrics, Division of Medical Genetics, Center for Genomic and Computational Biology, Duke University, Durham, North Carolina 27708, USA; Department of Integrative Biology and Pharmacology, McGovern Medical School, Houston, Texas 77030, USA

**Keywords:** Crohn’s disease, ulcerative colitis, cis-regulatory, NR2A1, NR2A2, gnotobiotic, Fiaf, H3K4me1, H3K27ac, DNase hypersensitivity, microbiome

## Abstract

Microbiota influence diverse aspects of intestinal epithelial physiology and disease in part by controlling tissue-specific transcription of host genes. However, host genomic mechanisms mediating microbial control of host gene expression are poorly understood. Using an unbiased screening strategy, we found that the zebrafish Hepatic nuclear factor 4 alpha (Hnf4a) transcription factor specifically binds and activates a microbiota-suppressed intestinal epithelial transcriptional enhancer. Genetic analysis disclosed that zebrafish *hnf4a* activates nearly half of the genes that are suppressed by microbiota, suggesting microbiota negatively regulate Hnf4a. In support, analysis of genomic architecture in mouse intestinal epithelial cells revealed that microbiota colonization leads to activation or inactivation of hundreds of enhancers along with drastic genome-wide reduction of Hnf4a and Hnf4g occupancy. Interspecies meta-analysis suggests Hnf4a may mediate microbial contributions to inflammatory bowel disease pathogenesis. These results indicate Hnf4a plays a critical conserved role in maintaining intestinal homeostasis in response to microbiota and inflammation.

## INTRODUCTION

All animals face the fundamental challenge of building and maintaining diverse tissues while remaining sensitive and responsive to their environment. This is most salient in the intestinal epithelium which performs important roles in nutrient absorption and barrier function while being constantly exposed to complex microbial communities (microbiota) and nutrients within the intestinal lumen. The presence and composition of microbiota in the intestinal lumen influence diverse aspects of intestinal development and physiology including dietary nutrient metabolism and absorption, intestinal epithelial renewal, and edification of the host immune system. Abnormal host-microbiota interactions are strongly implicated in the pathogenesis of inflammatory bowel diseases (IBD), including Crohn’s disease (CD) and ulcerative colitis (UC) (Sartor and Wu 2016). Studies in mouse and zebrafish models of IBD have established that impaired intestinal epithelial cell (IEC) responses to microbiota are a key aspect of disease progression (Bates et al. 2007; Kamada et al. 2013; Marjoram et al. 2015). Improved understanding of the molecular mechanisms by which microbiota evoke host responses in the intestinal epithelium can be expected to lead to new strategies for preventing or treating IBD and other microbiota-associated diseases.

The ability of IEC to maintain their physiologic functions and respond appropriately to microbial stimuli is facilitated through regulation of gene transcription. Genome-wide comparison of transcript levels in intestinal tissue or isolated IEC from mice reared in the absence of microbes (germ-free or GF) to those colonized with a microbiota (conventionalized or CV) have revealed hundreds of genes that have significantly increased or decreased mRNA levels following microbiota colonization (Camp et al. 2014). Interestingly, many mouse genes that are transcriptionally regulated by microbiota have zebrafish homologs that are similarly responsive, suggesting the existence of evolutionarily-conserved regulatory mechanisms (Rawls et al. 2004). For example, the protein hormone Angiopoetin-like 4 (Angptl4, also called Fiaf) is encoded by a single ortholog in the mouse and zebrafish genomes, and microbiota colonization results in significant reductions in transcript levels in the small intestinal epithelium of both host species (Bäckhed et al. 2004; Camp et al. 2012). Whereas these impacts of microbiota on host IEC transcriptomes and their downstream consequences have been extensively documented, the upstream transcriptional regulatory mechanisms remain poorly understood.

Specification and tuning of gene transcription proceeds in part through interactions between transcription factors (TFs) and their sequence-specific binding to *cis*-regulatory DNA. *Cis*-regulatory regions (CRRs) harbor binding sites for multiple activating or repressing TFs and are generally associated with nucleosome depletion and specific post-translational modifications of histone proteins within adjacent nucleosomes when acting as poised (H3K4me1) or active (H3K27ac) enhancers (Creyghton et al. 2010). Antibiotic administration can impact transcript levels and histone modifications in IECs (Thaiss et al. 2016), however it’s unclear if these changes are indirect effects caused by alterations to microbiota composition, direct effects of the antibiotic on host cells, or by the effects of remaining antibiotic-resistant microbiota (Morgun et al. 2015). Previous studies have shown that histone deacetylase 3 is required in IECs to maintain intestinal homeostasis in the presence of microbiota (Alenghat et al. 2013), and that overall histone acetylation and methylation in the intestine is altered by microbiota colonization (Krautkramer et al. 2016). However, the direct and specific effects of the microbiota on host CRRs and subsequent transcriptional responses in IECs remain unknown.

Our previous work predicted key roles for one or more nuclear receptor TFs in microbial down regulation of IEC gene expression (Camp et al. 2014), but the specific TF(s) were not identified. Nuclear receptors are ideal candidate TFs for integrating microbe-derived signals, since for many their transcriptional activity can be positively or negatively regulated by the binding of metabolic or hormonal ligands (Evans and Mangelsdorf 2014). However, the roles of nuclear receptors in host responses remain poorly understood, and no previous study has defined the impact of microbiota on nuclear receptor DNA binding. Nuclear receptors are a metazoan innovation. The earliest animals encoded a single nuclear receptor orthologous to Hepatocyte nuclear factor 4 (Hnf4; nuclear receptor subfamily NR2A) (Bridgham et al. 2010). Despite subsequent duplication and diversification, distinct Hnf4 TFs remain encoded in extant animals including mammals (Hnf4a, Hnf4g) and fishes (Hnf4a, Hnf4b, Hnf4g) (Supplemental Fig. S1G). Hnf4a serves particularly important roles in IECs, where it binds CRRs and activates expression of genes involved in IEC maturation and function (Stegmann et al. 2006). IEC-specific knockout of mouse *Hnf4a* results in spontaneous intestinal inflammation similar to human IBD (Darsigny et al. 2009). In accord, genetic variants at human *HNF4A* are associated with IBD as well as colon cancer (Marcil et al. 2012; Chellappa et al. 2016). Hnf4a is predicted to bind a majority of IBD-linked CRRs and regulate IBD-linked genes (Haberman et al. 2014; Meddens et al. 2016). Similarly, genetic variants near human *HNF4G* have been associated with obesity and IBD (Franke et al. 2007; Berndt et al. 2013). Importantly, these diverse roles for Hnf4 TFs in host physiology have only been studied in host animals colonized with microbiota. Therefore, the role of Hnf4 in host-microbiota interactions and the implications for human IBD remain unknown.

## RESULTS

### *hnf4a* is essential for transcriptional activity from a microbiota-suppressed cis-regulatory DNA region

To identify transcriptional regulatory mechanisms underlying microbial control of host gene expression, we took advantage of a previously identified microbiota-responsive CRR termed in3.4 located within the third intron of zebrafish *angptl4* (Fig. 1A). A GFP reporter construct under control of in3.4 termed *in3.4:cfos:gfp* drives tissue specific expression of GFP in zebrafish IEC and is suppressed by microbiota colonization, recapitulating the microbial suppression of zebrafish *angptl4* (Camp et al. 2012). However, the factor(s) that mediate microbial suppression of in3.4 were unknown. Using a yeast one-hybrid (Y1H) assay, we tested the capacity of 150 TFs expressed in the zebrafish digestive system to bind in3.4 (Supplemental Fig. S1A,B; Supplemental Table S1) and detected an interaction only with *hnf4a, hnf4b,* and *hnf4g* (Fig. 1B). When either of two predicted Hnf4 motifs in in3.4 are mutated, the Hnf4-in3.4 interaction in the Y1H assay and intestinal GFP expression in *in3.4:cfos:gfp* zebrafish is strongly reduced (Supplemental Fig. S1C-F). Interestingly, while *gata4, gata5,* and *gata6* have predicted motifs in in3.4 (Camp et al. 2012) these TFs did not interact in the Y1H assay. This suggests that HNF4 TFs are capable of binding in3.4 directly and HNF4 binding sites are necessary for directing in3.4-based transcription *in vitro* and in the intestine.

**FIGURE 1.**
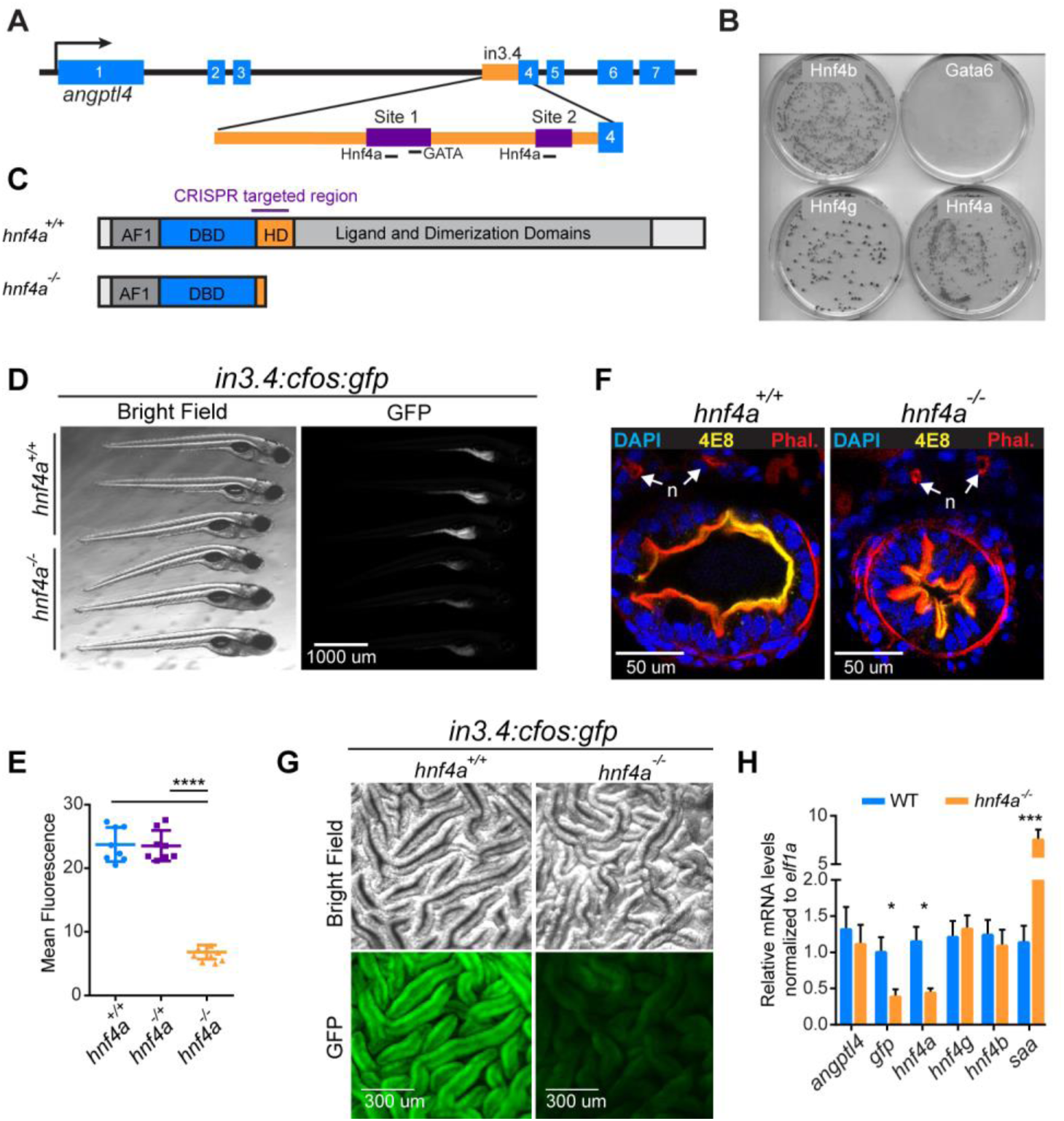
Zebrafish *hnf4a* is required for robust *in3.4:cfos:gfp* activity. (A) Schematic of the microbiota-suppressed zebrafish enhancer, in3.4, highlighting the regions required for intestinal activity (purple) which both contain putative Hnf4 binding sites (Site 1 and Site 2) (Camp et al. 2012). (B) Image of 4 plates from the Y1H assay showing the *hnf4* family of transcription factors capable of binding in3.4 and driving expression of the antibiotic resistance reporter gene. (C) Hnf4a^+/+^ and Hnf4a^−/−^ protein cartoons showing the DNA binding domain (DBD) and hinge domain (HD). We characterized the two with the largest lesions, a -41 deletion in the hinge domain and a +25 insertion in the hinge domain which both result in frame-shift early-stop codons and significantly reduced transcript. (D) Stereofluorescence GFP and bright field microscopy showing representative *hnf4a*^+/+^ (top 3) and *hnf4a*^−/−^ (bottom 3) 6dpf *in3.4:cfos:gfp* zebrafish. Genotype was blinded and samples were arranged by intensity of GFP fluorescence. (E) GFP fluorescence (mean ± sem) in *hnf4a*^+/+^ (n = 8), *hnf4a*^+/−^ (n = 8) and *hnf4a*^−/−^ (n = 8) 6dpf *in3.4:cfos:gfp* zebrafish (Two-tailed t-test: t = 17.84, 16.51, respectively, df = 14, and **** p < 0.0001). (F) Confocal microscopy showing representative axial cross sections in 6dpf *hnf4a*^+/+^ (n = 4) and *hnf4a*^−41/−41^ (n = 4) larval zebrafish. 4e8 antibody (yellow) labels the intestinal brush border, DAPI (blue) and phalloidin (red), and nephros (n). (G) Bright field microscopy (top) and stereofluorescence GFP (bottom) for representative *hnf4a*^+/+^ (n = 3) (left) and *hnf4a*^−/−^ (n = 3) (right) dissected intestinal folds from adult *in3.4:cfos:gfp* zebrafish. (H) Relative mRNA levels (mean ± sem) in *hnf4a*^+/+^ (n = 3) and *hnf4a*^−/−^ (n = 3) adult zebrafish intestinal epithelial cell as measured by qRT-PCR. Two-tailed t-test: t = 0.93, 5.22, 6.56, 10.65, 0.75, 0.94 respectively, df = 4, and * p <0.05, *** p < 0.001). See also Supplemental Figures S1 and S2.

We hypothesized that the *hnf4* transcription factor family is required to mediate microbial suppression of in3.4 activity. Although the Y1H assay demonstrated all 3 zebrafish Hnf4 members are capable of binding in3.4, we concentrated our efforts on understanding the function of *hnf4a* because it is the most highly conserved Hnf4 family member (Supplemental Fig. S1G; Supplemental Table S10) and has well-documented roles in intestinal physiology (San Roman et al. 2015). To that end, we generated *hnf4a* mutant zebrafish using the CRISPR/Cas9 system (Fig. 1C; Supplemental Fig. S2A-C,E). Unlike mouse whole-animal *Hnf4a* knockout animals that fail to initiate visceral endoderm development and die during early embryogenesis (Duncan et al. 1997), zebrafish *hnf4a* mutant animals are viable and survive to sexual maturity (Supplemental Fig. S2D) providing new opportunities to study the roles of Hnf4a in host-microbiota interactions.

To determine if *hnf4a* is essential for in3.4 activity, we crossed mutant *hnf4a* alleles into the *in3.4:cfos:gfp* transgenic reporter line. GFP expression was significantly reduced in the absence of *hnf4a* suggesting that *hnf4a* activates in3.4 (Fig. 1D,E,G,H). This loss of GFP expression in *hnf4a*^−/−^ mutants was not associated with overt defects in brush border development or epithelial polarity in larval stages (Fig. 1F), nor in the establishment of intestinal folds during adult stages (Fig. 1G). However, intestinal lumen of mutant larvae was reduced in size at 6 days post fertilization (dpf) compared to WT siblings (Fig. 1F; Supplemental Fig. S2F). Together, these data indicate *hnf4a* is essential for robust activity of a microbiota-suppressed CRR. Unlike *in3.4:cfos:gfp, angptl4* is expressed in multiple tissues and cell types (Camp et al. 2012). To determine if intestinal *angptl4* expression is dependent on *hnf4a* function, we isolated RNA from IECs from *hnf4a*^+/+^ and *hnf4*^−/−^ adult *in3.4:cfos:gfp* zebrafish and performed qRT-PCR. Adult IECs (AIECs) from *hnf4a*^−/−^ have significant reductions in mRNA for *gfp, fabp2* (a known Hnf4a target in human cell lines) (Klapper et al. 2007), and *hnf4a* compared to *hnf4a*^+/+^ controls. However, *angptl4* expression remained unchanged in *hnf4*^−/−^ AIECs compared to WT, suggesting *angptl4* transcript levels in the adult intestine are regulated by additional mechanisms and not solely from in3.4 or Hnf4a (Fig. 1H). Transcript levels for *hnf4g* and *hnf4b* in *hnf4a*^−/−^AIEC were also unchanged. Together, these results establish that Hnf4a is required for in3.4 activity in IECs and raises the possibility that Hnf4a may have broader roles in mediating host transcriptional and physiological responses to microbiota.

### Hnf4a activates transcription of genes that are suppressed upon microbiota colonization

To better define the roles of *hnf4a* in microbiota response and other aspects of digestive physiology, we used RNA-seq to compare mRNA levels from digestive tracts isolated from *hnf4a*^+/+^ and *hnf4a*^−/−^ zebrafish larvae in the presence (CV) or absence of a microbiota (GF; Fig. 2A). Consistent with our previous studies(Rawls et al. 2004; Kanther et al. 2011), comparison of wildtype zebrafish reared under CV vs GF conditions revealed differential expression of 598 genes that were enriched for processes such as DNA replication, oxidation reduction, and response to bacterium (Fig. 2B,D; Supplemental Fig. S3D; Supplemental Tables S2, S4). Strikingly, disruption of the *hnf4a* gene caused gross dysregulation of the transcriptional response to microbiota colonization, with the total number of microbiota responsive genes (CV vs GF) increasing to 2,217. Furthermore, comparison of the *hnf4a* mutant (Mut) vs wild type (WT) genotypes revealed differential expression of many genes in the CV condition (2,741 genes) and GF condition (1,441 genes) that inform a general role for Hnf4a in regulating genes in the intestinal tract (Fig. 2D,E). Principal components analysis (Supplemental Fig. S3A) and hierarchical clustering (Fig. 2B) of FPKM values indicated that *hnf4a* genotype had a complex contribution to regulating genes involved in both response to the microbiota and digestive physiology.

**FIGURE 2.**
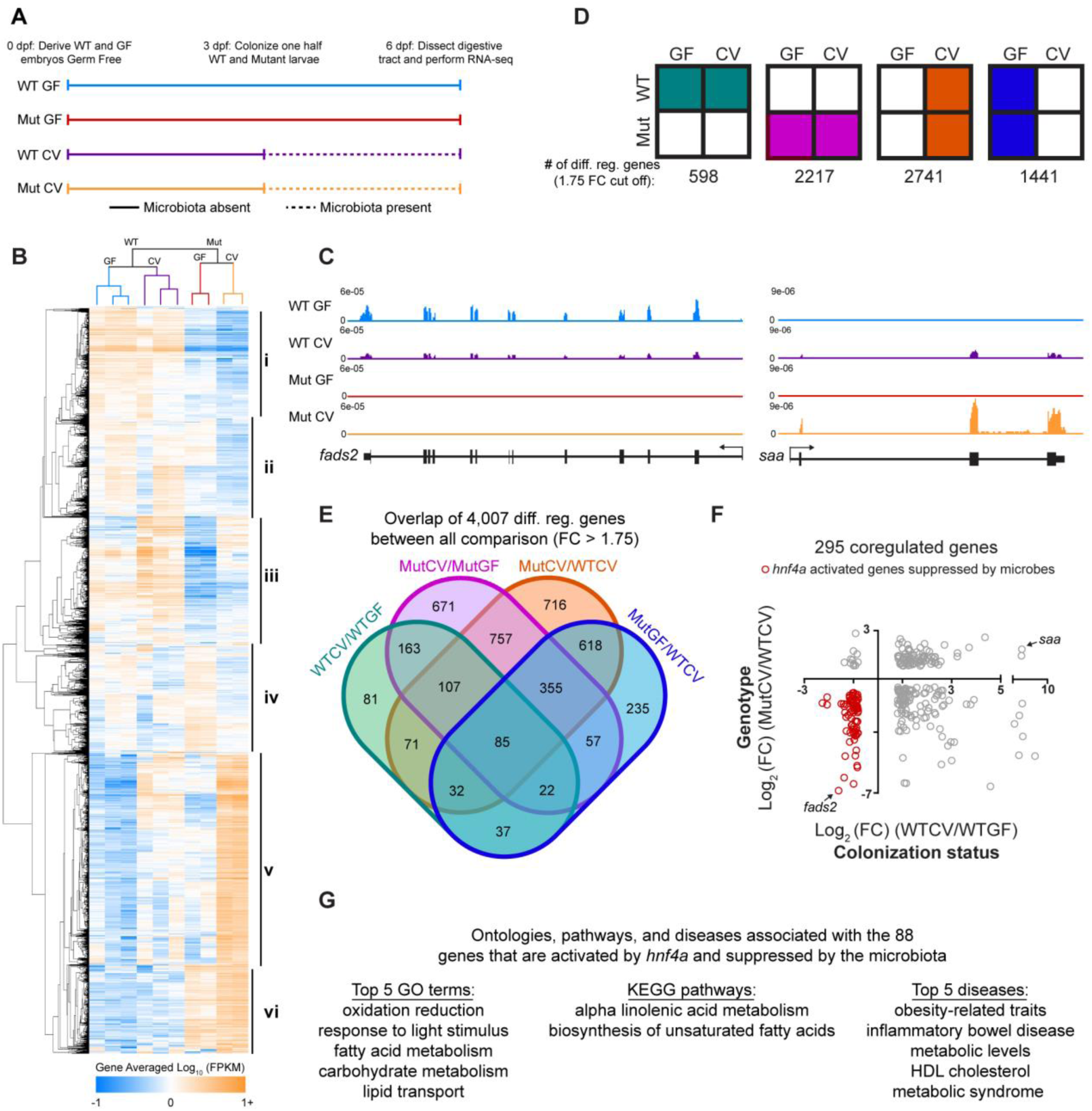
Hnf4a activates the majority of coregulated genes that are suppressed by the microbiota. (A) Schematic showing the experimental timeline for zebrafish digestive tract GF and CV *hnf4a*^+/+^ and *hnf4a*^−/−^ RNA-seq experiment. (B) Hierarchical relatedness tree and heatmap of differentially regulated genes in mutant and gnotobiotic zebrafish digestive tracts. Gene averaged log_10_ FPKMs for the biological replicates are represented for each of the 4,007 differentially regulated genes. (C) Representative RNA-seq signal tracks at *fatty acid-desaturase 2 (fads2), serum amyloid a (saa)* loci. (D) Summary of the total number of differentially expressed genes between indicated conditions (GF and CV) and genotype (WT and *hnf4a*^−/−^ (Mut)). (E) 4-way Venn diagram showing overlaps between all 4,007 differentially regulated genes. (F) The 295 coregulated genes were plotted using the log_2_ (FC) calculated in the WTGF/WTCV comparison (X-axis) and WTCV/MutCV (Y-axis). The 88 out of 98 genes that are activated by *hnf4a* but suppressed by the microbiota are highlighted (red) and (G) their GO term, KEGG pathway and disease associations are listed. See also Supplemental Figure S3.

Because we found that *hnf4a* activates the microbiota-suppressed intestinal CRR, in3.4, we hypothesized that this may represent a general regulatory paradigm for other microbiota-influenced CRRs and genes across the genome. When we compared the 598 genes that were microbiota responsive in wildtype digestive tracts with the 2,741 genes that *hnf4a* regulates in CV digestive tracts we found these lists shared 295 genes that included *fads2* and saa, both of which have human orthologs that are either implicated *(FADS1/2)* or markers (*SAA*) of IBD (Plevy et al. 2013; Costea et al. 2014) (Fig. 2C-F). While loss of Hnf4a could be pleiotropic, strikingly, the overlap between these subsets reveals that a disproportionate 88 of the 98 (~90%) microbiota-suppressed genes are activated by *hnf4a* (Fig. 2F; Supplemental Table S2). These 88 genes represent almost half of all 185 genes suppressed by the microbiota. These data suggest, like its role at in3.4, *hnf4a* plays a critical role in directly activating a large percentage of genes that are suppressed by microbial colonization. Interestingly, despite a clear enrichment of metabolism themed ontologies and pathways, we found IBD was among the top 2 diseases associated with genes that are activated by Hnf4a but suppressed by the microbiota (Fig. 2G; Supplemental Table S11). Based on these results, we hypothesized that Hnf4a DNA binding is lost upon microbial colonization within CRRs associated with microbiota-suppressed genes.

### Hnf4a binding sites are enriched in promoters near genes associated with microbiota-regulated H3K27ac marks

Previous attempts to identify microbial responsive enhancers genome-wide were complicated by the lack of significant changes in chromatin DNase accessibility between GF and CV IECs from mouse colon and ileum (Camp et al. 2014). These previous findings suggested other chromatin dynamics may be involved in regulating the IEC response to microbiota. We therefore sought to provide a genomic context for understanding how the microbiota alter Hnf4a activity and chromatin modifications in IECs by performing RNA-seq, DNase-seq, and ChIP-seq for the enhancer histone modifications H3K4me1 and H3K27ac, and the Hnf4 TF family members Hnf4g and Hnf4a in CV and GF conditions totaling 35 datasets. We conducted these experiments in jejunal IEC from gnotobiotic mice because: (1) ChIP-grade antibodies for mouse Hnf4a and Hnf4g are available, (2) the larger organ size provided sufficient numbers of IECs for ChIP-seq experiments, and (3) we speculated that the roles of Hnf4a in host response to microbiota may be conserved to mammals. We first performed DNase-seq in jejunal IEC from mice reared GF or colonized for two weeks with a conventional mouse microbiota (CV) to determine the impact of microbiota colonization on chromatin accessibility (Fig. 3A). In accord with previous studies that that tested for chromatin accessibility in ileal or colonic IECs from GF or CV mice (Camp et al. 2014), we similarly found no differential DNase hypersensitivity sites (DHSs) in GF or CV jejunum (data not shown, but see Supplemental Fig. S4A; Supplemental Tables S6, S8). These data indicate that gross accessibility changes in chromatin do not underlie the transcription of microbiota-responsive genes in IECs.

**FIGURE 3.**
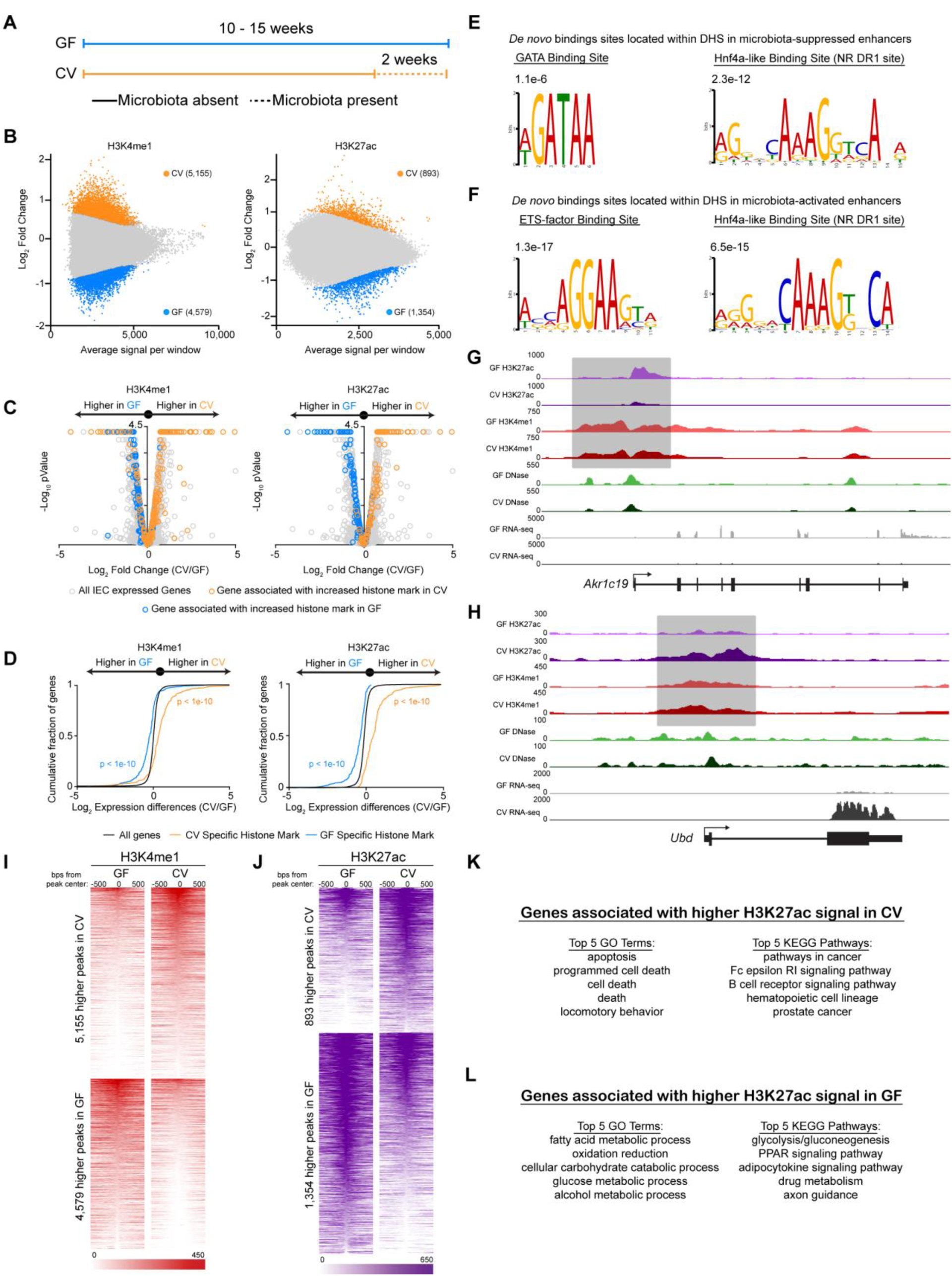
Microbiota selectively induce enhancer activity near genes that are upregulated upon microbiota colonization. (A) Schematic showing the gnotobiotic experimental timeline for testing mRNA levels and chromatin architecture in GF and CV. (B) MA plots from DEseq2 analysis (FDR < 0.01) of H3K4me1 (n = 3 per condition) (left) and H3K27ac (n = 2 per condition) (right) ChIP-seq from GF and CV mouse jejunal IECs. Colored dots signify regions significantly enriched for a histone mark in GF (blue) or CV (orange). We found 4,579 unique H3K4me1 and 1,354 unique H3K27ac peaks in GF and 5,155 unique H3K4me1 and 893 unique H3K27ac peaks in CV. (C) Volcano plots showing pairwise comparison of RNA expression between GF and CV jejunal IECs. Blue and orange dots represent genes associated with a region enriched for H3K4me1 (left) or H3K27ac (right) signal in GF or CV. (D) Two-sided Kolmogorov-Smirnov goodness-of-fit test shows a positive relationship on average between the presence of a region enriched for H3K4me1/H3K27ac signal in a specific colonization state and increased transcript abundance of a neighboring gene in that same colonization state. (E) Top *de novo* binding site motifs found in DHSs that are flanked by regions enriched with H3K27ac signal in GF (E) or CV (F). Representative ChIP-seq tracks highlighting a microbiota-regulated gene associated with differential histone marks in GF (G) (*Akr1c19*, *Aldo-keto reductase 1c19)* or CV (H) (*Ubd, Ubiquitin D*). Heatmaps showing the average GF and CV H3K4me1 (I) or H3K27ac (J) signal at the 1000 bps flanking differential sites. (K-L) GO terms and KEGG pathways enriched in genes associated with differential H3K27ac sites shown in J. See also Supplemental Figure S4.

To test if other metrics of chromatin utilization were dynamically regulated by microbiota, we performed ChIP-seq from GF and CV mouse jejunal IECs for histone marks H3K4me1 and H3K27ac that are enriched at poised enhancers and active enhancers, respectively (Fig. 3B). By determining the single-nearest gene TSS within 10kb of the differential histone marks and overlaying these data with our new RNA-seq datasets, we found that regions that gain poised (H3K4me1) and activated (H3K27ac) enhancers upon colonization are associated with genes that have increased transcript levels upon colonization (Fig. 3C,H-K; Supplemental Fig. S4I; Supplemental Tables S3, S6, S8). Similarly, regions that lose poised and active enhancers upon colonization are associated with microbiota-suppressed genes (Fig. 3C,G,I,J,L; Supplemental Fig. S4J; Supplemental Tables S3, S6, S8). A two-sided Kolmogorov-Smirnov goodness-of-fit test shows a positive relationship between differential H3K4me1/H3K27ac region and increased transcript abundance of nearby genes in the same colonization state (Fig. 3D). Collectively, we identified for the first time a genome-wide map of hundreds of newly identified microbial regulated CRRs, suggesting that microbiota regulation of host genes is mechanistically linked to histone modifications changes more than gross chromatin accessibility changes (Camp et al. 2014).

We leveraged this novel atlas of microbiota-regulated enhancers and accessible chromatin to determine which TFs are predicted to bind to these regions. An unbiased analysis found that Hnf4a binding site motifs were enriched in promoters of genes associated with microbiota-suppressed enhancers (Supplemental Fig. S4E), and STAT1 binding site motifs were enriched in promoters of genes associated with microbiota-activated enhancers (Supplemental Fig. S4F). Interestingly, DHS sites associated with differentially active enhancers were enriched for two different sets of TF binding sites. DHSs flanked by microbiota-inactivated enhancers were enriched for nuclear receptor DR1 sites and GATA binding sites (Fig. 3E). DHS sites associated with microbiota-activated enhancers were similarly enriched for the nuclear receptor DR1 binding sites but also for STAT/IRF-like and ETS binding sites (Fig. 3F). These data suggest that nuclear receptors like Hnf4a may play a central role in IEC responses to microbial colonization.

### Microbiota colonization is associated with a reduction in Hnf4a and Hnf4g cistrome occupancy

To directly evaluate the impact of microbiota on Hnf4a activity, we tested the plasticity of the genome wide distribution of Hnf4s in response to microbial colonization. Hnf4a bound 28,901 and Hnf4g bound 21,875 across the genome in GF conditions in jejunal IECs with ~80% of these sites being bound by both TFs. In striking contrast, the number of sites bound by Hnf4a and Hnf4g in CV conditions was ~10 fold less (Fig. 4A,B; Supplemental Tables S5, S8). Of the 3,964 Hnf4a binding sites detected in CV there were only 267 Hnf4a sites that were specific to the CV condition (Supplemental Fig. S5A,C; Supplemental Table S8). Yet, the genes associated with these Hnf4a sites that are retained in CV are enriched for ontologies and pathways fundamental to intestinal epithelial biology (Supplemental Fig. S5B). However, we did find that the average CV Hnf4a signal was significantly increased at Hnf4a sites associated with microbiota-induced genes relative to those Hnf4a sites associated with microbiota-suppressed genes, suggesting Hnf4a may play a limited role in genes upregulated by colonization (Supplemental Fig. S5F). In contrast, GF Hnf4a ChIP signal was equivalent at Hnf4a sites associated with microbiota-suppressed and induced genes (Supplemental Fig. S5F). We do not believe that the reduction of Hnf4a binding is the result of chromatin quality in a particular condition since there are genomic locations where GF and CV Hnf4a sites appeared to have equivalent signal (Fig. 4C). Furthermore, ChIP enrichment in these IEC preparations for another zinc finger TF, CTCF, was unaffected by microbiota colonization (Supplemental Fig. S5D). This indicates that the observed reduction of Hnf4 ChIP signal in CV IECs is a result of microbiota on Hnf4 binding, and is not the result of altered ChIP efficiency or sample quality in the different conditions. The dramatic loss of Hnf4a and Hnf4g DNA binding upon colonization is consistent with Hnf4a acting as a potent activator of microbiota-suppressed genes.

**FIGURE 4.**
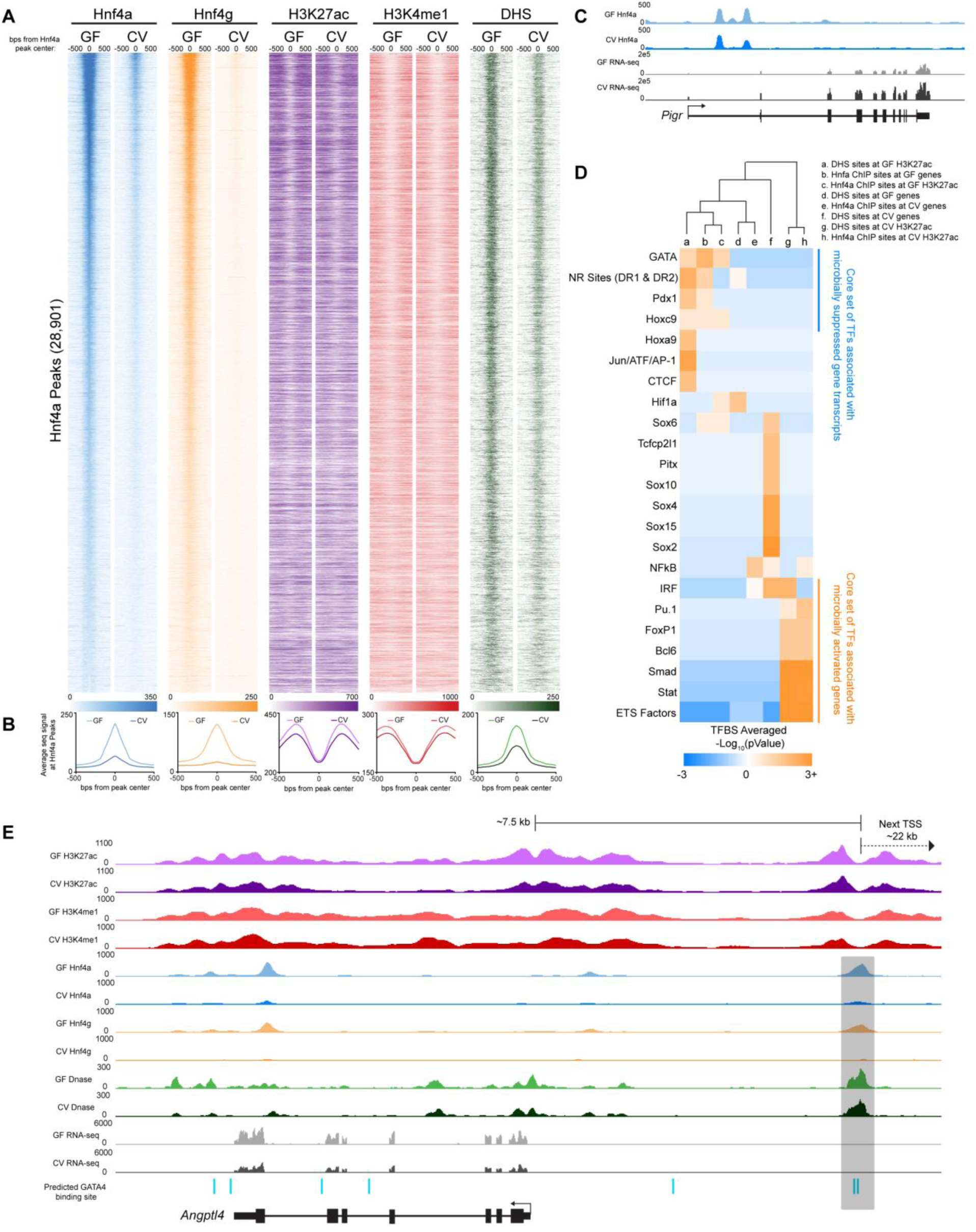
Microbiota colonization results in extensive loss of Hnf4a and Hnf4g DNA binding in IEC. (A) Heatmaps showing the average GF and CV ChIP-seq or DNase-seq signal at the 1000 bps flanking Hnf4a sites found in GF.(B) Line plots showing the average GF (light-colored line) and CV (dark-colored line) ChIP-seq and DNase-seq RPKM-normalized signal for the indicated TF, histone mark or DHS at the 1000 bps flanking Hnf4a sites found in GF (Hnf4a: n = 3 per condition; Hnf4g: n = 4 per condition; H3K27ac n = 2 per condition; H3K4me1: n = 3 per condition; DNase: N = 3 for CV, n = 2 for GF). (C) Representative signal tracks highlighting a microbiota-induced gene (*Pigr, Polymeric immunoglobulin receptor*) that is associated with an Hnf4a peak with similar signal in both GF and CV jejunal IECs. (D) Heatmap showing the enrichment of TFBS motifs within 50 bps of the DHS or Hnf4a peak maxima. (E) Representative signal track at *Angptl4* highlighting two GATA4 sites within an Hnf4a bound region. See also Supplemental Figure S5.

We further speculated that certain coregulatory sequence-specific transcription factors may also contribute to regulating transcription with Hnf4 at these sites. To explore this possibility, we searched for TF motifs associated with Hnf4a ChIP sites and found an enrichment of putative binding sites for TFs known to be involved in small intestinal physiology (GATA and Hoxc9) as well as nutrient metabolism (Pdx1) at both Hnf4a bound regions associated with genes and enhancers suppressed by microbes (Fig. 4D). We similarly found GATA sites located within an Hnf4a-bound CRR near murine *Angptl4* (Fig. 4E), similar to the coincident Hnf4 and GATA motifs in in3.4 (Camp et al. 2012). Furthermore, binding sites for TFs known to be involved in cell proliferation and cell death (ETS transcription factor family) are enriched near Hnf4a bound regions that intersect microbiota-induced enhancers (Fig. 4D). Collectively our integrative analyses of these novel ChIP-seq, DNase-seq, and RNA-seq datasets identifies a core set of putative microbiota-responsive TFs that may interact with Hnf4a to mediate microbial control of IEC gene expression. These results suggest Hnf4a plays a major role in integrating microbial signals to regulate gene expression, and raise the possibility that this novel microbiota-Hnf4a axis might contribute to human disease.

### Microbiota-mediated suppression of Hnf4a may contribute to IBD pathogenesis

Both Hnf4a and the intestinal microbiota have been separately implicated in the pathogenesis of inflammatory bowel diseases such as Crohn’s disease (CD) and ulcerative colitis (UC) (Ahn et al. 2008; Sartor and Wu 2016). However, a mechanistic link between microbiota and Hnf4a in the context of IBD pathogenesis has not been established. Previous transcriptomic studies have identified genes differentially expressed in ileal (iCD) and colonic CD (cCD) and UC (Arijs et al. 2009; Haberman et al. 2014) biopsies. We queried these human gene lists to identify one-to-one orthologs in mice, and referenced them against our new gnotobiotic mouse jejunal Hnf4a ChIP-seq data (Fig. 5A). Strikingly, the majority of human genes downregulated in all of these IBD datasets have mouse orthologs that are associated with an Hnf4a-bound region (Fig. 5B,C; Supplemental Table S7). Focusing on the iCD dataset from the largest of these previous studies (Haberman et al. 2014), we found differential iCD genes associated with Hnf4a sites are enriched for distinct ontologies and pathways that are dysregulated in IBD (Fig. 5H-K). In contrast to IBD, analysis of a transcriptomic dataset from human necrotizing enterocolitis (NEC) (Tremblay et al. 2016) revealed significantly less Hnf4a-bound regions near genes both induced and suppressed in NEC (Fig. 5B,C). These data indicate microbiota-dependent and microbiota-independent suppression of Hnf4a activity may play an important role in IBD pathologies, but not NEC.

**FIGURE 5.**
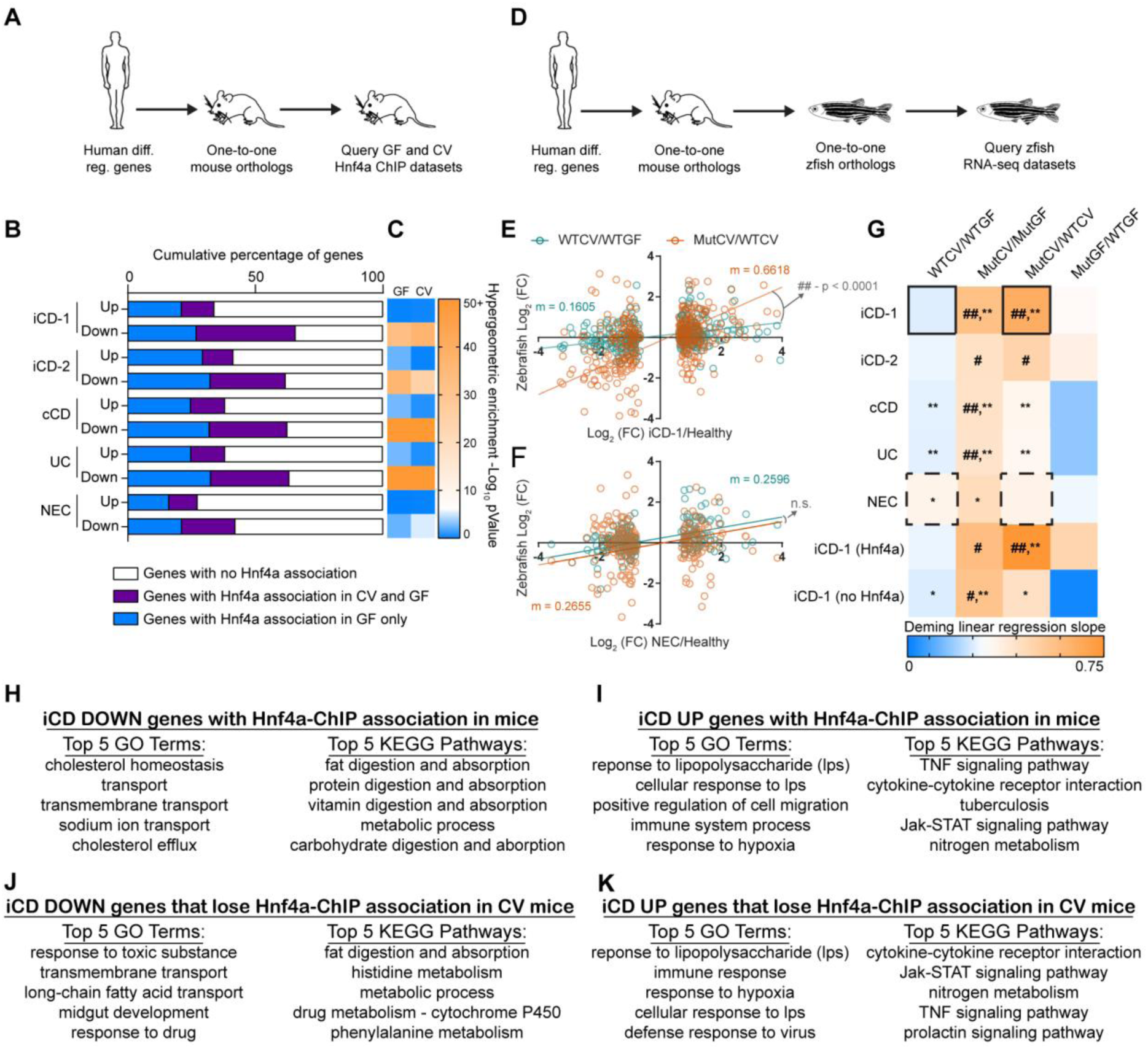
Microbiota suppression of Hnf4a activity is highly correlated with genes and intestinal processes suppressed in human IBD and conserved in zebrafish. (A) Flow chart showing the experimental design and filters used to identify IBD or NEC gene orthologs associated with mouse Hnf4a ChIP sites. (B) Bar chart showing the proportion of Hnf4a associations in GF and CV mouse jejunal IECs near human-to-mouse one-to-one gene orthologs differentially regulated in human pediatric ileal Crohn’s Disease (iCD-1), adult iCD (iCD-2), adult colonic Crohn’s Disease (cCD), adult ulcerative colitis (UC), or neonatal necrotizing enterocolitis (NEC). (C) Heatmap representing the -Log_10_ (pValue) of the enrichment of GF or CV Hnf4a-associated genes that are differentially regulated genes in the indicated IBD datasets. pVlaues were calculated using a hypergeometric enrichment analysis and converting all Hnf4a ChIP-associated mouse genes to human orthologs (GF = 5863 genes and CV = 2119 genes). (D) Flow chart showing the experimental design and filters used to identify correlations between gnotobiotic WT or mutant zebrafish gene expression and gene orthologs differentially expressed in human IBD or NEC. Because loss of *hnf4a* function in zebrafish appeared to more closely resemble iCD signature than cCD or UC, we performed pairwise comparisons of gene orthologs that are (1) differentially regulated in human iCD and (2) have a mouse Hnf4a ChIP association. Example of Deming linear regression analysis showing the correlation of Log_2_ (FC) between WTCV/WTGF (E) or MutCV/WTCV (F) zebrafish and pediatric iCD or NEC. m = slope of the line. (G) Heatmap representing slopes of Deming linear regression lines showing positive correlative relationships between the log_2_ gene expression fold changes of one-to-one orthologs from human diseases compared to log_2_ fold changes in zebrafish WTCV/WTGF, MutCV/MutGF, MutCV/WTCV, and MutGF/WTGF. Hash signs indicate slope of Deming linear regression lines is significantly greater than WTCV/WTGF comparison (#, p < 0.05; ##, p < 0.0001). Asterisks indicate slope of Deming linear regression line is significantly greater than MutGF/WTGF (*, p <0.05; **, p <0.001). Solid boxes correspond to slope of lines in panel 5D, and dashed boxes correspond to slope of lines in panel 5E. (HK) The top 5 GO terms and the top 5 KEGG pathways for indicated gene lists.

To assess if microbiota suppression of Hnf4a activity regulates genes differentially expressed in IBD, we queried the published human IBD and NEC gene expression datasets to identify human-mouse-zebrafish one-to-one-to-one orthologs that were differentially expressed in our RNA-seq analysis of gnotobiotic zebrafish *hnf4a* mutants (Fig. 5D). We found ortholog expression fold changes in human IBD/healthy comparisons most closely resembled the expression fold changes of MutCV/MutGF and MutCV/WTCV (Fig. 5E-G). Neither the WTCV/WTGF nor the MutGF/WTGF comparisons faithfully recapitulate the expression profiles of IBD/healthy comparisons. This indicates that both the microbiota and loss of *hnf4a* function are necessary to induce an IBD-like gene expression profile in zebrafish. Strikingly, the positive correlation and significant resemblance to the iCD-like gene signatures in the colonized *hnf4a*^−/−^ compared to colonized *hnf4a*^+/+^ zebrafish digestive tracts become even stronger when we limited our analysis to one-to-one orthologs that have an association with an Hnf4a bound region in mouse IECs (Fig. 5G). Together, these results suggest that Hnf4a protects the host and maintains transcriptional homeostasis in the presence of a microbiota and protects against an IBD-like gene expression signature.

## DISCUSSION

Over the course of animal evolution, the intestinal epithelium has served as the primary barrier between animal hosts and the complex microbial communities they harbor. IECs maintain this barrier and perform their physiological roles in nutrient transport and metabolism through dynamic transcriptional programs. The regulatory mechanisms that orchestrate these transcriptional programs represent potential therapeutic targets for a variety of human intestinal diseases including IBD. Here we discovered that Hnf4a activity and its transcriptional network are suppressed by microbiota. Hnf4a is the oldest member of the nuclear receptor TF family (Bridgham et al. 2010), and our findings in fish and mammals suggest that microbial suppression of Hnf4a may be a conserved feature of IEC transcriptional programs present in the common ancestor.

Our results suggest important new links between Hnf4a and microbiota in the context of human IBD. IBD patients, particularly those suffering from Crohn’s disease, often present with decreased serum low-density lipoprotein levels and reduced total cholesterol levels compared to healthy individuals (Hrabovsky et al. 2009; Agouridis et al. 2011). These serum levels are consistent with reduced transcript levels for genes involved in intestinal absorption and transport of lipid and cholesterol in ileal and colonic biopsies from IBD patients (Arijs et al. 2009; Haberman et al. 2014). Transcription factors, including nuclear receptors like Hnf4a and Fxr, are known to regulate bile acid production, lipid mobilization and cholesterol absorption have already been implicated in IBD (Ahn et al. 2008; Nijmeijer et al. 2011). However, our work is the first to demonstrate a role for microbiota in suppressing Hnf4a that may ultimately affect IBD pathogenesis. Hnf4a has been shown to play key roles in anti-oxidative and anti-inflammatory defense mechanisms (Marcil et al. 2010) so aberrant microbial suppression could promote a chronic inflammatory state. Hnf4a target genes are downregulated in human IBD (Arijs et al. 2009; Haberman et al. 2014) and mouse experimental colitis, and the Hnf4a target ApoA1 has been shown to be protective against intestinal inflammation in mice (Gkouskou et al. 2016). We speculate that the genes governed by this novel microbiota-Hnf4a axis may include additional anti- and pro-inflammatory factors that could provide new targets for IBD therapy.

Our results reveal similar effects of microbiota colonization and experimental colitis on Hnf4a cistrome occupancy, but the underlying molecular mechanisms are unresolved. DSS induced colitis results in reduced Hnf4a protein levels and altered cellular localization (Chahar et al. 2014), however our results indicate the microbiota neither reduce Hnf4a protein levels nor impact its nuclear localization in jejunal IECs two weeks after colonization (Supplemental Fig. S5H,I). Colonization of GF mice with microbiota initiates a transcriptional adaptation in the intestine that progresses for several weeks before reaching homeostasis (El Aidy et al. 2012), so further studies are needed to determine if Hnf4a activity is similarly altered at other stages of this adaptive process. However, these data collectively suggest that microbiota suppress Hnf4a activity in the jejunum through mechanisms distinct from those utilized in experimental colitis.

Despite the importance of Hnf4a in digestive and metabolic health (Palanker et al. 2009; Frochot et al. 2012), relatively little insight into its regulation in a biological context has been reported. *In vitro* and cell culture studies have identified possible suppressors and activators of Hnf4a including phosphorylation by AMPK and acetylation by CBP, both of which have been shown to induce Hnf4a activity (Soutoglou et al. 2000; Hong et al. 2003). Other studies have shown that Hnf4a regulation is controlled by alternative promoters which generate different isoforms(Huang et al. 2009). However, we did not detect differential *Hnf4a* exon usage by DEXseq (Li et al. 2015) in GF and CV mouse jejunal IECs (data not shown). Another facet of Hnf4a biology that has not been adequately explored is the identity of its endogenous ligand(s). Although historically considered an orphan nuclear receptor, several fatty acids including linoleic acid have been identified as ligands for Hnf4a (Hertz et al. 1998; Palanker et al. 2009; Yuan et al. 2009). Fatty acids are an attractive class of putative regulators of Hnf4a since the microbiota are known to augment FA absorption in zebrafish enterocytes (Semova et al. 2012).

Identification of Hnf4a ligands or other co-regulatory factors that mediate microbial control of Hnf4a could provide novel therapeutic targets for IBD (Mbodji et al. 2013).

In our attempt to understand how the microbiota regulate Hnf4a activity and host gene transcription, we were motivated to investigate if microbiota impact histone modification and chromatin accessibility in the mouse jejunum. Our findings support the model that microbiota alter IEC gene expression by affecting TF binding and histone modification at tissue-defined open chromatin sites (Camp et al. 2014). We provide the genomic addresses of hundreds of microbiota-regulated enhancers as well as the genes associated with these enhancers and Hnf4a binding sites. Similar to other findings in intraepithelial lymphocytes (Semenkovich et al. 2016), our work demonstrates a clear microbial contribution to the modification of the histone landscape in IECs and provides another important layer of regulation that orchestrate microbiota regulation of host genes involved in intestinal physiology and human disease. We were also able to establish a link between microbiota-regulated genes and enhancers and NR binding sites. These NR binding sites are coincident with a core set of TFs that are enriched near microbiota-suppressed enhancers/genes (GATA) or induced enhancers/genes (ETS-factors and IRF) (Supplemental Fig. S6). Coregulation by other TFs represents one possible mode of Hnf4a regulation by which the microbiota could suppress Hnf4a activity without impacting the gene transcription of all Hnf4a-associated genes.

## METHODS

### Yeast 1-Hybrid ORFeome Screen

The yeast 1-hybrid ORFeome screen was performed using the Clontech MatchmakerTM Gold Yeast One-hybrid Library Screening System (cat. 630491) protocol with the following exceptions: The Y1HGold yeast strain was transformed using standard yeast transformation procedures with BstBI digested pBait-AbAi containing either the WT or a SDM in3.4 or the p53 binding site sequence, and positive transformants were selected on SD/-URA media. In addition, a ORFeome library consisting of 148 zebrafish transcription factors cloned from adult zebrafish liver (Supplemental Table S1) into pDEST22 prey vectors containing an N-terminal GAL4-activation domain was utilized (Boyle et al. 2016). For additional information, see Supplemental Methods.

### Zebrafish Transgenesis and Imaging

Co-injections of Tol2 SDM or WT in3.4:cfos:gfp plasmid and transposase mRNA were performed as described (Camp et al., 2012) with the following exceptions: 50 – 100 zebrafish embryos were injected at the 1–2 cell stage with approximately 69 pg of plasmid DNA at a DNA:transposase ratio of 1:2. Three mosaic patches within a given tissue of an imaged fish were quantified for mean fluorescence intensity and averaged. Statistical significance was analyzed using Kruskal-Wallis one-way analysis of variance and Dunn’s multiple comparison test using GraphPad Prism software. For additional information, see Supplemental Methods.

### Zebrafish Mutagenesis

Targeted gene deletion of the *hnf4a* gene was performed using CRISPR/Cas9 nuclease RNA-guided genome editing targeting the fourth exon of *hnf4a*. The guide RNA sequences were designed using “CRISPR Design Tool” (http://crispr.mit.edu/). Guide RNAs (Supplemental Table S9) were generated from BamHI (New England Biolabs R0136L) digested pT7-gRNA plasmid (a gift from Wenbiao Chen and available from Addgene: http://www.addgene.org/46759/) and by performing an *in vitro* transcription reaction using MEGAshortscript T7 kit (Ambion/Invitrogen AM1354). For additional information, see Supplemental Methods.

### Mouse IEC Isolation for DNAse, ChIP and RNA-seq

Mice were euthanized under CO_2_ and cervical dislocation and placed on a chilled wax dissection pad. The small intestine was removed from the mouse and the jejunum was excised from the duodenum and ileum. Duodenum was defined as the anterior 5 cm of the midgut and ileum was defined as posterior 6 cm of midgut as described (Camp, et al 2014). Adipose and vasculature were removed from the tissue. The jejunum was opened longitudinally along the length of the tissue, exposing the lumen and epithelial cell layer. Luminal debris was washed away from the epithelia with ice cold sterile PBS. The tissue was temporarily stored in 10 ml of ice cold sterile PBS with 1x Protease Inhibitor (Complete EDTA-Free, Roche 1187350001) and 10 uM Y-27632 (ROCK I inhibitor, Selleck Chemicals S1049) to inhibit spontaneous apoptosis. The jejunum was moved into a 15 ml conical tube containing 3 mM EDTA in PBS with 1x protease inhibitor and 10 uM Y-27632. The tissue was placed on a nutator in a cold room for 15 minutes. The jejunum was removed from the 3 mM EDTA and placed on an ice cold glass petri dish with PBS containing 1mM MgCl2 and 2 mM CaCl2 with protease inhibitors and 10 uM Y-27632. Villi were scraped off of the tissue using a sterile plastic micropipette and placed into a new 15 ml conical tube. The isolated IECs were then crosslinked for ChIP or used for DNAse-seq or RNA-seq. For additional information, see Supplemental Methods.

### Mouse Intestine Immunofluorescence and Western Blot

Mid-jejunal tissue was dissected and cleaned as in the IEC villi isolation above. The whole, splayed open tissue was pinned to 3% agarose and fixed in 4% PFA overnight with gentle agitation at 4°C. The fixed tissue was washed 4 times with PBS for 15 minutes. The tissue was then permeabilized in PBS with 0.5% Tween 20 for 1.5 hours at room temperature. Following permeabilization, the tissue was blocked in 5% donkey serum in PBS with 0.1% Tween 20 for 2 hours at room temperature, incubated in primary and secondary antibodies and imaged on a Leica SP8 confocal microscope. Western blots were performed on non-crosslinked IEC lysates (see below) using standard chemoluminescence Western blot protocols. The western blot shown in Supplemental Fig. S5H is a representative of two experiments. For additional information, see Supplemental Methods.

### Bioinformatic and Statistical analysis

Sample sizes for zebrafish experiments (noted in figure legends) were selected based on genotype availability and transgenesis efficiency. All sample collection was performed two or more times on independent days. For sequencing experiments, statistical calls for differential gene expression were made by Cuffdiff using parameters stated above. For the zebrafish RNA-seq experiment Next-Gen sequencing was performed once and at the same time to avoid batch effects: WTGF and WTCV (n = 3); MutGF and MutCV (n = 2). We originally collected n = 3 MutGF and MutCV biological replicates, however, using pre-established criteria and to avoid RNA contamination, we excluded one biological replicate from all analysis from these groups because of sequencing reads that mapped within the deleted *hnf4a* exon in the *hnf4a*^−/−^ genotype.

GF mice were randomly chosen by gnotobiotic staff for microbiota colonization (CV) based on their availability and litter sizes. All sample collection was performed two or more times per condition on independent days. GF and CV mouse samples were collected on different days. For sequencing experiments, statistical calls for differential gene expression and differential peak calls were made by Cuffdiff, MACS2, and DEseq2 using parameters stated above. For the mouse RNA-seq experiment Next-Gen sequencing was performed once and at the same time to avoid batch effects: GF (n = 2) and CV (n = 2). Paired GF and CV ChIP and library amplification was performed simultaneously. Typically, biological ChIP replicates were sequenced on different days and were always paired with the other condition (i.e. CV and GF were always sequenced together). The number of biological ChIP replicates (noted in figure legends) was dependent on reproducibility between ChIP samples and/or our ability to determine statistical differential sites using DEseq2 (for H3K4me1 and H3K27ac).

All statistical metrics (except where otherwise noted) were performed in Graphpad Prism 7.01. Deming linear regression was used for Fig. 5 because it is a stronger and more accurate assessment of correlation when both the x and y variables have experimental error. Details regarding the other statistical tests used in this study can be found in the figure legends or above.

For additional information, see Supplemental Methods.

## Dataset Accessions

Sequence datasets are available at GEO accession number GSE90462.

## ACKNOWLEDGEMENTS

We thank Balfour Sartor, Scott Magness, and Maureen Bower for assistance with gnotobiotic mice, and Wenbiao Chen and Stacy Horner for sharing reagents. We also thank the Genomic Sequencing Laboratory at HudsonAlpha Institute for Biotechnology and the Duke Sequencing and Genomic Technologies Facility. This work was supported by grants from the National Institutes of Health (R01-DK081426, U24-DK097748, P01-DK094779, R24-OD016761, and P30-DK34987).

